# Mapping protein conformational landscapes from crystallographic drug fragment screens

**DOI:** 10.1101/2024.07.29.605395

**Authors:** Ammaar A. Saeed, Margaret A. Klureza, Doeke R. Hekstra

## Abstract

Proteins are dynamic macromolecules. Knowledge of a protein’s thermally accessible conformations is critical to determining important transitions and designing therapeutics. Accessible conformations are highly constrained by a protein’s structure such that concerted structural changes due to external perturbations likely track intrinsic conformational transitions. These transitions can be thought of as paths through a conformational landscape. Crystallographic drug fragment screens are high-throughput perturbation experiments, in which thousands of crystals of a drug target are soaked with small-molecule drug precursors (fragments) and examined for fragment binding, mapping potential drug binding sites on the target protein. Here, we describe an open-source Python package, COLAV (COnformational LAndscape Visualization), to infer conformational landscapes from such large-scale crystallographic perturbation studies. We apply COLAV to drug fragment screens of two medically important systems: protein tyrosine phosphatase 1B (PTP-1B), which regulates insulin signaling, and the SARS CoV-2 Main Protease (MPro). With enough fragment-bound structures, we find that such drug screens also enable detailed mapping of proteins’ conformational landscapes.

## Introduction

While often shown as single structures, proteins exhibit dynamic behavior necessary for their function^1–3^, e.g. binding and releasing ligands^4^, modulating activity^5^, and reversibly shielding the active site^6^. Hence, proteins are better thought of as populating ensembles of structural states or conformations. Individual protein molecules transition frequently between these conformations through the concerted motions of their amino acids. For many proteins, there are only a handful of accessible backbone conformations at physiological temperatures, all separated by distinct concerted motions^2,7^.

Consequently, proteins can often be thought of as residing on a conformational landscape that describes metastable conformations and the concerted motions necessary to transition between them^8^. Ideally, conformational landscapes would be inferred from experimental structures and would succinctly recapitulate the known conformational diversity of a target protein. Additionally, these empirical landscapes would suggest thermally accessible concerted motions between conformations—probable temporal sequences of conformational change sometimes referred to as conformational reaction coordinates or transition paths^9–11^. Such conformational landscapes for validated protein drug targets would suggest particular conformations to (de)stabilize to enhance or inhibit functional activity. These conformations can then be targeted by the design of a small molecule that binds the drug target within the active site (orthosteric) or elsewhere (allosteric).

Existing biophysical methods can experimentally characterize aspects of a protein’s conformational landscape, e.g., by nuclear magnetic resonance (NMR) spectroscopy^12^, fluorescence resonance energy transfer spectroscopy^13^, electron paramagnetic resonance spectroscopy^14^, and room-temperature X-ray crystallography^6,15^. These techniques probe the equilibrium distribution of a desired conformational ensemble. However, such measurements generally reflect the ground state of the protein and only provide limited insight into the presence and/or nature of any alternate, higher-energy conformations. For large proteins and protein complexes, cryogenic electron microscopy (cryo-EM) and electron tomography (cryo-ET) can capture small populations of metastable conformations directly^16^, and machine learning methods are beginning to pave the way for the identification of these rare protein states^17,18^. Yet, determining high-resolution structures of metastable states through cryo-EM or cryo-ET remains an ongoing challenge, due to the need for a vast quantity of correctly classified particle images.

An alternative approach to studying these excited states is to directly perturb the protein of interest. These perturbations alter the conformational landscape, stabilizing otherwise short-lived excited states. Common methods to introduce perturbations include mutation of the protein and addition of substrate/transition-state analogs. Once the protein has been perturbed, the stabilized states can be examined via standard biophysical techniques. Though the efficacy of this approach has been demonstrated in a variety of model systems^19–21^, designing individual perturbations can be time-consuming and may only explore a limited portion of the conformational landscape.

An ideal approach to mapping protein conformational landscapes would be to subject the protein of interest to a large number of distinct perturbations that are just strong enough to bias the energetics of particular conformations by a few *k*_B_*T* and then determine the structure of the protein under each perturbation^22,23^. Crystallographic drug fragment screens constitute an intriguing approximation to this ideal experiment: in these high-throughput crystallographic screens, many crystals of the same drug target are each soaked with a unique drug fragment and are then subjected to the standard X-ray crystallography pipeline. Advances in automation at the Diamond Light Source^24^ and elsewhere, paired with novel data processing software^25,26^, have enabled these screens to solve thousands of protein structures within days, some of which contain bound drug fragments. Importantly, these drug fragment screens may yield information valuable for drug design beyond the immediate identification of drug fragment/binding site pairs: a comprehensive exploration of the protein’s conformational landscape.

To test this idea, we developed a software package known as COLAV (COnformational LAndscape Visualization) that calculates three different representations of protein structure— dihedral angles, pairwise distances, and strain—to quantify structural change across a group of crystal structures. COLAV is an open-source, Python-based software, freely available at https://github.com/Hekstra-Lab/colav. Using COLAV, we show that sets of crystal structures can be used to construct a map of a protein’s conformational landscape and infer correlated regions within the protein. We then ask whether the conformational landscape constructed from structures obtained only from a crystallographic drug fragment screen is consistent with a map of the landscape based on structures obtained using a variety of perturbations (e.g., mutants, substrate analogs, and inhibitors) available from the Protein Data Bank (PDB)^27^. We find that the drug fragment-derived map provides a partial view of the conformational landscape that is consistent with the landscape derived from the complete dataset. The drug fragment-derived map becomes substantially more complete with increasing scale of the crystallographic drug fragment screen.

## Methods

### Structural representations

We implemented three methods to represent a protein structure in COLAV: backbone dihedral angles (*ϕ*, *ω*, and *ψ*), pairwise distances between Cα atoms, and strain. We implemented these methods on top of the Scientific Python stack (NumPy^28^, SciPy^29^, and BioPandas^30^). Dihedral angles and distances were calculated according to standard methods, and strain was calculated according to previously published frameworks^31,32^ described briefly below. To ensure consistent features across each protein dataset, we truncated structures at the N and C termini and then removed any structures missing backbone atoms between the truncated endpoints. For PTP-1B, we calculated representations between residues 7 and 279 (inclusive). For “focused PCA” of the PTP-1B L16 loop, we only used representations between residues 236 and 244 (inclusive). For MPro, we calculated representations between residues 3 and 297 (inclusive). If alternate conformations had been modeled for any atoms, then we included only the “A” conformer in our calculations. In our strain implementation, we calculated three different variants of strain: strain tensor, shear tensor, and shear energy. We used the off-diagonal elements of the shear tensor as inputs for principal component analysis (PCA). Use of COLAV is illustrated in the accompanying Jupyter Notebooks available at https://github.com/Hekstra-Lab/colav.

### Data analysis

We analyzed these structural representations using the Scikit-Learn implementation of PCA, using 10 principal components (PCs) and otherwise default parameters^33^. Because of the inherent periodicity present in dihedral angles, we linearized these features by calculating the sine and cosine of each angle and using the resulting tuple as the input feature for PCA. To determine a per-residue measure of importance for each method (“residue contributions”), we transformed the coefficients of the principal components as follows. For dihedral angles, we first summed the absolute values of the sine and cosine coefficients of the same dihedral angle to determine a per-angle, per-residue measure. We also summed the absolute values of these per-angle measures into a single per-residue measure. For the pairwise distance representation, we summed the absolute value of all coefficients pertaining to each residue. For the strain-based representation, we summed the absolute value of the off-diagonal elements of the shear matrix for each residue.

We also analyzed these structural representations using the Scikit-Learn implementation of t-distributed Stochastic Network Embedding^34^ (t-SNE) and the Umap-Learn implementation of Uniform Manifold Approximation and Projection^35^ (UMAP). We initialized both of these latter methods randomly; we did not observe major differences in the clustering of structures when using different seeds. To identify groupings of structures similar to each other in the MPro dataset, we used the Scikit-Learn implementation of the *k*-means algorithm with default settings^33^. In our assessment of the role of dataset size, we generated MPro datasets of varying size by sampling the complete MPro dataset (without replacement) each time.

To establish the coupling between regions of PTP-1B, we performed Fisher exact tests for independence (https://www.socscistatistics.com/tests/). This test asserts as a null hypothesis that the variables used are independent and as an alternative hypothesis that there is a dependence structure among the variables. We tested for conditional independence by adding the chi-square statistics of two two-way tests and comparison to the null distribution (chi-square with two degrees of freedom) as described in Ch. 5, “Analysis of Discrete Data”, (https://online.stat.psu.edu/stat504/book/).

### Dataset construction

For PTP-1B, we retrieved 165 structures of the human enzyme from the Protein Data Bank (PDB) in March 2022 with a sequence identity of 90% or higher compared to wild-type PTP-1B. We also retrieved 187 structures of PTP-1B bound to fragment ligands from a crystallographic drug fragment screen^36^ that were identified either by Pan-Dataset Density Analysis (PanDDA)^25^ alone or after tandem processing by cluster4x^26^ and PanDDA. We retrieved all PTP-1B files in the PDB file format (hereafter .pdb).

For MPro, we retrieved all 1,830 crystallographic drug fragment screen structures in March 2022 from the Fragalysis database^37–41^. We retrieved all 1,015 other MPro structures from the PDB in July 2023. We excluded MPro structures from an ensemble refinement study of MPro at multiple temperatures (7MHL, 7MHM, 7MHN, 7MHO, 7MHP, 7MHQ)^42^; these temperature-induced effects dominated the analysis, masking the native conformational landscape of MPro. Several MPro structures were too large to download in the .pdb format, so we downloaded them in the mmCIF file format. We subsequently converted them to the .pdb format using an online GEMMI tool^43^.

Before feature extraction, we aligned structures of PTP-1B or MPro using THESEUS v3.3.0^44^, as superposing structures of the same protein was crucial for proper strain calculations. Where noted, we also idealized the backbone dihedral angles of each structure separately using Representation of Protein Entities (RoPE)^45^.

## Results and Discussion

### A framework for examining conformational change

COLAV offers three different structural representations to summarize differences between conformations, each with a distinct emphasis (Table S1 summarizes the functions available in COLAV). Dihedral angles and pairwise distances are internal coordinates, meaning that they are measures calculated from atomic coordinates regardless of the orientation of the protein. Therefore, these calculations can be performed on individual structures and do not require alignment of protein structures. Dihedral angles efficiently summarize local backbone dynamics of individual residues or loops by capturing these motions in only a few features, while pairwise distances better capture global protein dynamics, such as breathing motions^6^.

In contrast, strain analysis is a directional measure of the structural deformations accompanying conformational transitions. Using the strain analysis framework of previous studies^31,32^, all the structures must be aligned and compared to a designated reference structure. Here, the notion of continuous strain is discretized, instead focusing on individual atoms and their surrounding atomic neighborhoods—nearby atoms within 8 Å. By comparing the atomic neighborhoods in the working and reference structures, discrete analogs to continuous strain can be estimated, which then describe directional deformations of the desired structure relative to the reference. Notably, strain measurements pick up on regions with relative motion, for example around hinge points, while ignoring rigid-body-like motion, e.g., within subdomains.

### COLAV representations distinguish between known PTP-1B conformations

We applied all three methods implemented in COLAV to infer the conformational landscape of protein tyrosine phosphatase 1B (PTP-1B) from crystal structures. PTP-1B is a validated drug target for type II diabetes^46^ and breast cancer^46,47^, and has been implicated in Alzheimer’s disease^48^. Although there has been major pharmacological interest in PTP-1B, no drugs targeting PTP-1B have successfully made it through stage II clinical trials^49^. One major reason is that the PTP-1B active site is highly conserved across the protein tyrosine phosphatase family, making it difficult to design competitive inhibitors without off-target effects *in vivo*^50,51^. The PTP-1B active site is also charged, limiting the effective availability of charged competitive inhibitors that must cross a cell’s plasma membrane^51^. For these reasons, there has been widespread interest in allosterically targeting and modulating PTP-1B activity^52^. It is of particular interest, then, to discover surface sites allosterically coupled with the active site^36,53,54^.

To do so, we first analyzed a set of 352 crystal structures of PTP-1B obtained from the PDB (165 individual structures and 187 structures from a drug fragment screen performed by Keedy *et al*.^36^). Using principal component analysis (PCA), we found that each structural representation of conformational change implemented in COLAV separated the conformations into the same four clusters of distinct, known conformations (Fig. 1). These four conformations are described by the conformational states of the WPD and L16 loops (WPD loop/L16 loop): open/open (Fig. 1a top-left), open/closed (Fig. 1b bottom-left), closed/open (Fig. 1c top-right), and closed/closed (Fig. 1d bottom-right). For dihedral angles and strain, the first two PCs clustered these conformations (Fig. 1a, c); for pairwise distances, the first and third PCs clustered these conformations (Fig. 1b; PC2 determines regions with large motions relative to the rest of PTP-1B). We also applied two non-linear dimensionality reduction methods, t-distributed stochastic network embedding (t-SNE) and uniform manifold approximation and projection (UMAP), to the structural representations. These methods similarly clustered PTP-1B structures (Fig. S1), indicating that the PCA clusters were representative of the major groupings in the PTP-1B structures. We next asked whether inconsistent refinement practices for the deposited structures and/or deviations from ideal geometry in individual structures could explain the observed structural heterogeneity. To examine this possibility, we repeated the analysis after applying Representation of Protein Entities (RoPE)^45^ to all the PTP-1B structures to idealize and standardize the bond distances and bond angles across the dataset. In RoPE, the backbone dihedral angles of the structures are adjusted to match the original atomic coordinates. PCA identified the same PTP-1B clusters after pre-processing the data (Fig. S2a, b, e), confirming that individual refinement artifacts did not meaningfully affect the results.

**Figure 1:**
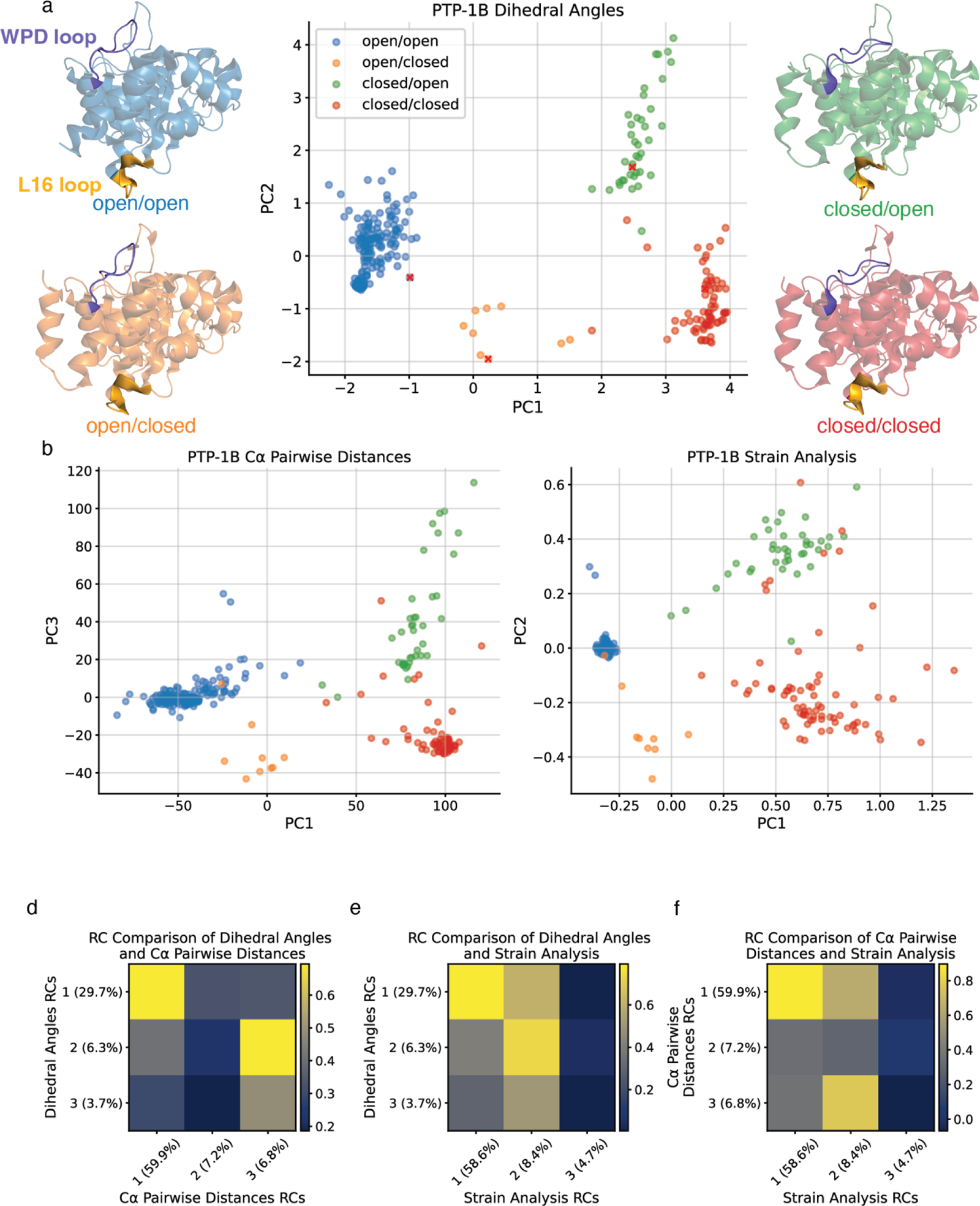
Conformational landscape of PTP-1B inferred using three different structural representations and colored by conformation. (a) PTP-1B conformational landscape by dihedral angles, flanked by representative PTP-1B structures of the four major conformations labeled by the conformational state of the WPD loop (purple) and L16 loop (yellow): (open/open: 1NWL, open/closed: 4QBW, closed/open: 1PXH, closed/closed: 1SUG). (b) PTP-1B conformational landscape by Cα pairwise distances; note that PC3 is shown on the *y*-axis. (c) PTP-1B conformational landscape by strain analysis. (d-f) Correlation coefficient matrix comparing RCs 1-3 for (d) dihedral angles and Cα pairwise distances; (e) dihedral angles and strain; (f) Cα pairwise distances and strain.

The three different structural representations implemented in COLAV can each capture different aspects of conformational change. It is conceivable that local conformational changes take place without much global change and are therefore primarily detectable by monitoring dihedral angles. Another possibility is that global change can be related to only a few dihedral angles, e.g., in hinge motion, but be detectable elsewhere as changing distances to other parts of the protein. Lastly, it is possible that coupled conformational changes are separated by regions of almost imperceptible change—possibly a common case for proteins^55–57^. To compare the conformational changes revealed by each representation, we calculated residue contributions (RCs) from the coefficients of each of the principal components (PCs), combining per residue the contributions of the sines and cosines of the dihedral angles (for the dihedral angle representation), of distances to all other residues (for the Cα pairwise distance representation), or off-diagonal components of the shear matrix (for the strain representation), respectively, as described in the Methods. By calculating the correlation between these RCs for each pair of representations (Figs. 1d-f, S3), we find that the residue contributions underlying PC1 and PC2 (“RC1” and “RC2”) for dihedral angles and for strain are strongly correlated (0.79 comparing RC1s and 0.74 comparing RC2s), respectively. Both RC1 and RC2 of these two representations show a correlation with the residue contributions underlying PC1 and PC3 for pairwise distances (Fig. 1d,f). As expected, however, the residue contributions are not perfectly correlated, indicating differences in the aspects of conformational change captured by each representation.

The PCs distinguish conformational clusters by the states of the WPD loop (Fig. 2a, b; residues 176-188) and L16 loop (Fig. 2a-c; residues 237-243). The active-site WPD loop participates in the PTP-1B catalytic mechanism, while the L16 loop is located ∼15 Å away (Fig. 1a). Both loops can take on open and closed states, and all four possible combinations of their states are present in the existing crystal structures. These loops account for most of the conformational heterogeneity present in the PTP-1B dataset (dihedral angles: 36.1% of total variance captured by the first two principal components, Cα pairwise distances: 66.6%, and strain analysis: 67.0%). In the WPD loop-open state, the loop is positioned such that the active-site pocket is exposed, facilitating substrate access and product release (Fig. 1a-left). In the WPD loop-closed state, the loop binds the substrate and covers the active site pocket, facilitating catalysis^4^ (Fig. 1a-right). The L16 loop states differ most saliently by the position of lysine 239 (K239)^36^. In the open state, the sidechain atoms of K239 interact primarily with the solvent (Fig. 1a-top). In the closed state, the sidechain atoms of K239 interact with other residues in the protein (Fig. 1a-bottom). By distinguishing the states of the WPD and L16 loops, PCA captures the major conformational heterogeneity present in crystal structures of PTP-1B.

**Figure 2:**
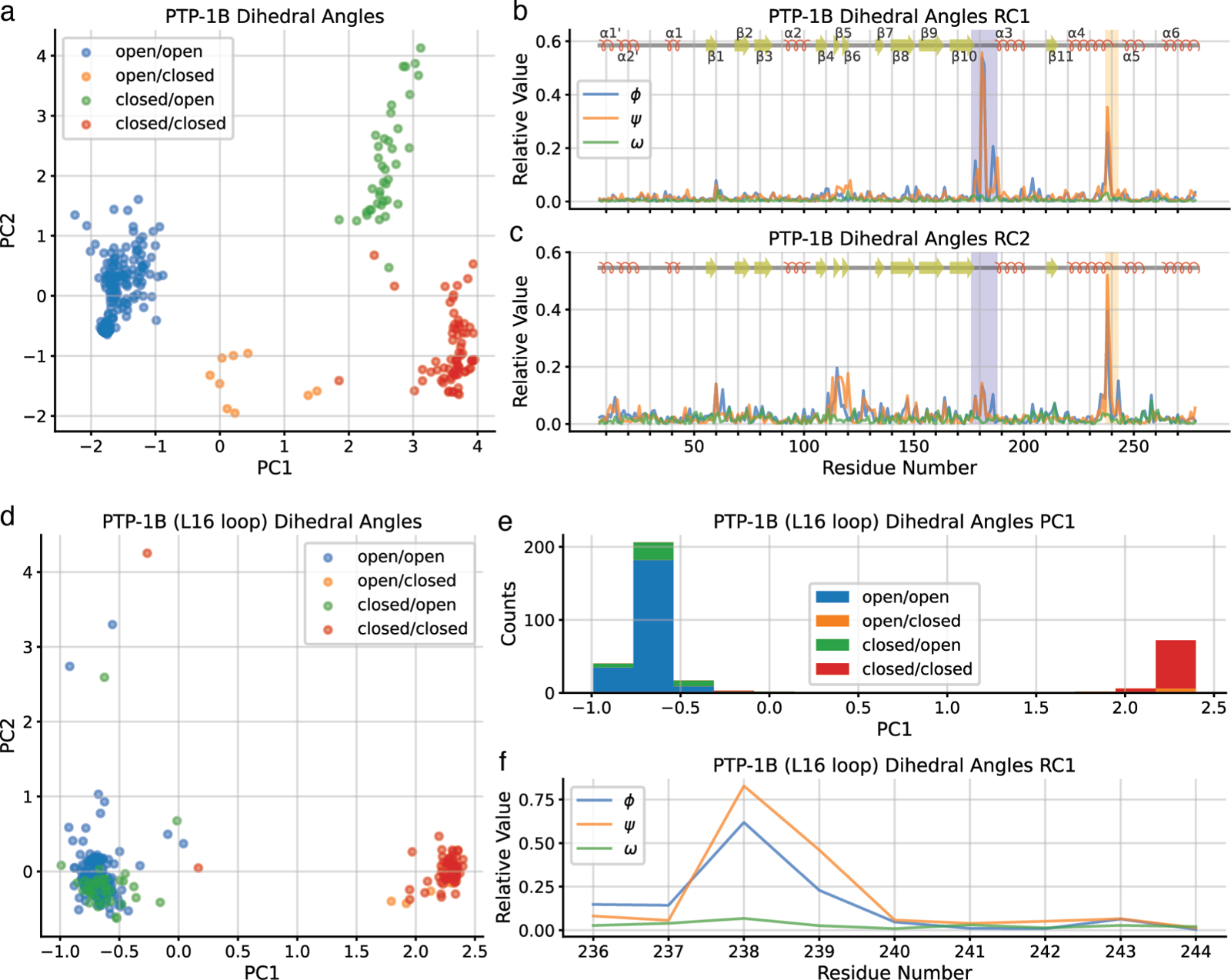
The dihedral angle representation distinguishes between conformations of PTP-1B based on the conformations of the WPD loop and L16 loop. (a) PTP-1B conformational landscape by dihedral angles by PC1 and PC2. (b) Residue contributions to principal component 1 (PC1), with WPD loop (residues 176-188) indicated by a purple box and L16 loop (residues 237-243) in a yellow box. (c) Residue contributions to PC2, with WPD loop in purple box and L16 loop in yellow box. (d) PTP-1B L16 loop conformational landscape by dihedral angles colored by conformation. (e) Histogram of PTP-1B structures according to PC1 of the focused PCA. (f) Residue contributions to PC1 of the focused PCA.

Could this conformational clustering be caused by crystal packing interactions, rather than the effects of perturbations introduced in individual structures? The most common space group of PTP-1B crystals in our dataset is the P3_1_21 space group, with 293 structures. The space groups of other PTP-1B crystals are P2_1_2_1_2_1_ (29), P12_1_1 (9), C121 (9), P3_2_21 (7), and P4_1_2_1_2 (2). As we show in Figure S4, the set of structures from crystals in the P3_1_21, P2_1_2_1_2_1_, and P12_1_1 space groups each encompasses all four major conformational clusters. Only the two structures from crystals in the P4_1_2_1_2 space group take on only a single conformation (closed/open). Since PTP-1B molecules across diverse space groups adopted different conformations, we conclude that crystal packing artifacts cannot account for the conformational clusters highlighted by PCA. Instead, these crystal structures represent semi-random samples from the PTP-1B conformational landscape.

### COLAV enables detection of correlated regions in PTP-1B

Although the crystal structures deposited in the PDB for any protein do not, together, constitute a valid thermodynamic ensemble, there is a long history of interpreting frequencies observed in crystal structures in thermodynamic terms^58–61^, most recently extending to the interpretation of AlphaFold parameters in energetic terms^62,63^. In this spirit, the statistical correlations observed as principal components can be interpreted as (rough) energetic couplings. Since the conformational landscapes determined by PCA were equivalent for all structural representations, we focus here on the dihedral angle representation (Fig. 1a). We interpreted the first principal component (PC), which accounts for 29.7% of the total variance, to indicate a coupling between the WPD loop and L16 loop (Fig. 2b). Indeed, previous experimental studies using multi-temperature X-ray crystallography^36^ and NMR^53,54^ have strongly suggested that these two loops are allosterically coupled. We interpreted the second PC, which accounts for 6.3% of the variance, to indicate additional motion of the L16 loop independent of the WPD loop (Fig. 2c). This observation suggests two possibilities. Either the L16 loop undergoes two distinct motions—one coupled to the WPD loop and another decoupled from the WPD loop—or the L16 loop undergoes a single motion that is not always coupled to the WPD loop. To differentiate between these possibilities, we performed a focused PCA on the dihedral angles of the L16 loop (Fig. 2d). We find that the L16 loop has a single dominant motion (Fig. 2d-f) that distinguishes between the open and closed states of the loop; this motion accounts for 63.5% of the variance in this focused PCA. Thus, the L16 loop undergoes a single motion that is not always coupled to the WPD loop.

To examine this coupling more closely, we considered the confounding effect of the C-terminal α7 helix, which has previously been implicated in allosteric coupling within PTP-1B^53^ and forms contacts with both loops in their respective closed states. We had initially excluded the α7 helix from our analysis to avoid missing values, as the α7 helix can transition between an ordered, folded helix state and a disordered state that is not crystallographically observable. However, we noticed that the α7 helix typically takes on the ordered state when at least one of the WPD or L16 loops takes on their respective closed conformations (Table 1). We hypothesized that the exclusion of the α7 helix had led to the observed inconsistencies in the coupling of the two loops. Within the PTP-1B dataset, we find that the presence of an ordered α7 helix greatly increases the probability of finding the closed state of each loop (∼40x for the L16 loop and ∼50x for the WPD loop). This suggests a cooperative mechanism in which binding of a ligand or inhibitor in the active site can drive concerted loop closure and ordering of the α7 helix.

**Table 1:** Assessing the correlations of the WPD loop, L16 loop, and α7 helix through χ^2^ test of independence. Contingency table comparing PTP-1B conformations of the WPD loop, the L16 loop, and the α7 helix. Calculated p-values are based on a Fisher exact test.

|  | Open<br>L16 | Closed<br>L16 | total |
| --- | --- | --- | --- |
| Open<br>WPD | **221** | 4 | 225 |
| Closed<br>WPD | 23 | 1 | 24 |
| **total** | 244 | 6 | 249 |

P-value: 0.40

**Table 1: Assessing the correlations of the WPD loop, L16 loop, and $\alpha 7$ helix through $\chi^2$ test of independence.**
|  | Open<br>L16 | Closed<br>L16 | total |
| --- | --- | --- | --- |
| Open<br>WPD | 7 | 5 | 12 |
| Closed<br>WPD | 16 | **72** | 88 |
| **total** | 23 | 77 | 100 |

P-value: 0.006

**Table 2:**
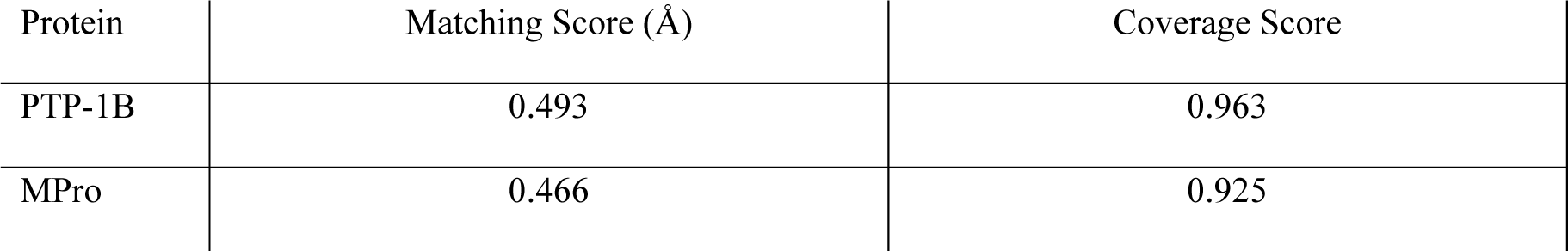
Matching and coverage scores comparing PDB-only and fragment screen-only structures for PTP-1B and MPro. The matching score reports on the similarity of the datasets by RMSD, and a smaller score implies that the datasets are more similar. The coverage score reports on diversity of structures between datasets, and the highest score of 1 implies that the datasets are similarly diverse.

| Protein | Matching Score ($\text{\AA}$) | Coverage Score |
| --- | --- | --- |
| PTP-1B | 0.493 | 0.963 |
| MPro | 0.466 | 0.925 |

To formally test for a coupling between the three regions of PTP-1B, we performed a three-way chi-square test of independence (Table 1; treating structures as independent observations), finding strong evidence that these regions are not independent (*p* ∼ 10^−158^). To assess the role of the α7 helix, we next tested how the correlation between the states of the WPD loop and L16 loop depends on the state of this helix (by Fisher’s exact test). Given a disordered α7 helix, we find no significant evidence for coupling of the WPD and L16 loops (however, the L16 loop is rarely in the closed state when the α7 helix is disordered, limiting the power of this test). Given an ordered α7 helix, the states of the two loops are strongly coupled to each other (*p* = 0.006; Fisher’s exact test). We can, in addition, reject the hypothesis that the state of the α7 helix solely specifies the state of each loop, as the loop states are not conditionally independent given the state of the α7 helix (*p* = 0.005; chi-squared test). Moreover, ligands are not necessary for the protein to visit states with closed WPD and L16 loops and an ordered α7 helix. For instance, apo structures collected at temperatures above 100 K (6B8E, 6B8T, 6B8X) show electron densities consistent with both states at each of these regions. In addition, several mutations can stabilize apo PTP-1B with the WPD and L16 loops in their closed states and an ordered α7 helix (1PA1, 6OLQ, 6OMY, 6PFW, 7KEN). The two loops are therefore coupled to each other and to the α7 helix, although the exact molecular mechanism remains unclear.

Detailed analysis of the COLAV results further showed active site deformation consistent with oxidation of the active-site catalytic cysteine residue Cys215 (Fig. 3a,b). Oxidation dynamics of this residue play a critical role in its function^64–67^ through a self-regulatory mechanism in PTP-1B^65^ and (when fully oxidized) degradation (Fig. 3a,b)^68^. The most striking of several oxidized states is a cyclized state in which a sulphenyl-amide bond between the Sγ atom of Cys215 and the backbone nitrogen atom of Ser216 forms a five-membered ring. Structures of oxidized conformations (1OEM and 1OES) show deformations at active site loops, matching RC4 (accounted for 3.5% of total variance) and RC5 (accounted for 2.9% of total variance) of the dihedral angle representation (Fig. 3d,e). Only six PTP-1B structures present in the dataset (∼2%) have oxidized cysteine states modeled, and PCA distinguishes these structures from structures in the native, reduced state (Fig. 3c, top-right corner). However, it is possible that low levels of oxidation in PTP-1B crystals are present more widely in the structures^36^, impacting the average electron density and, therefore, structure coordinates. Overall, applying PCA to COLAV results successfully identified these rare conformations.

**Figure 3:**
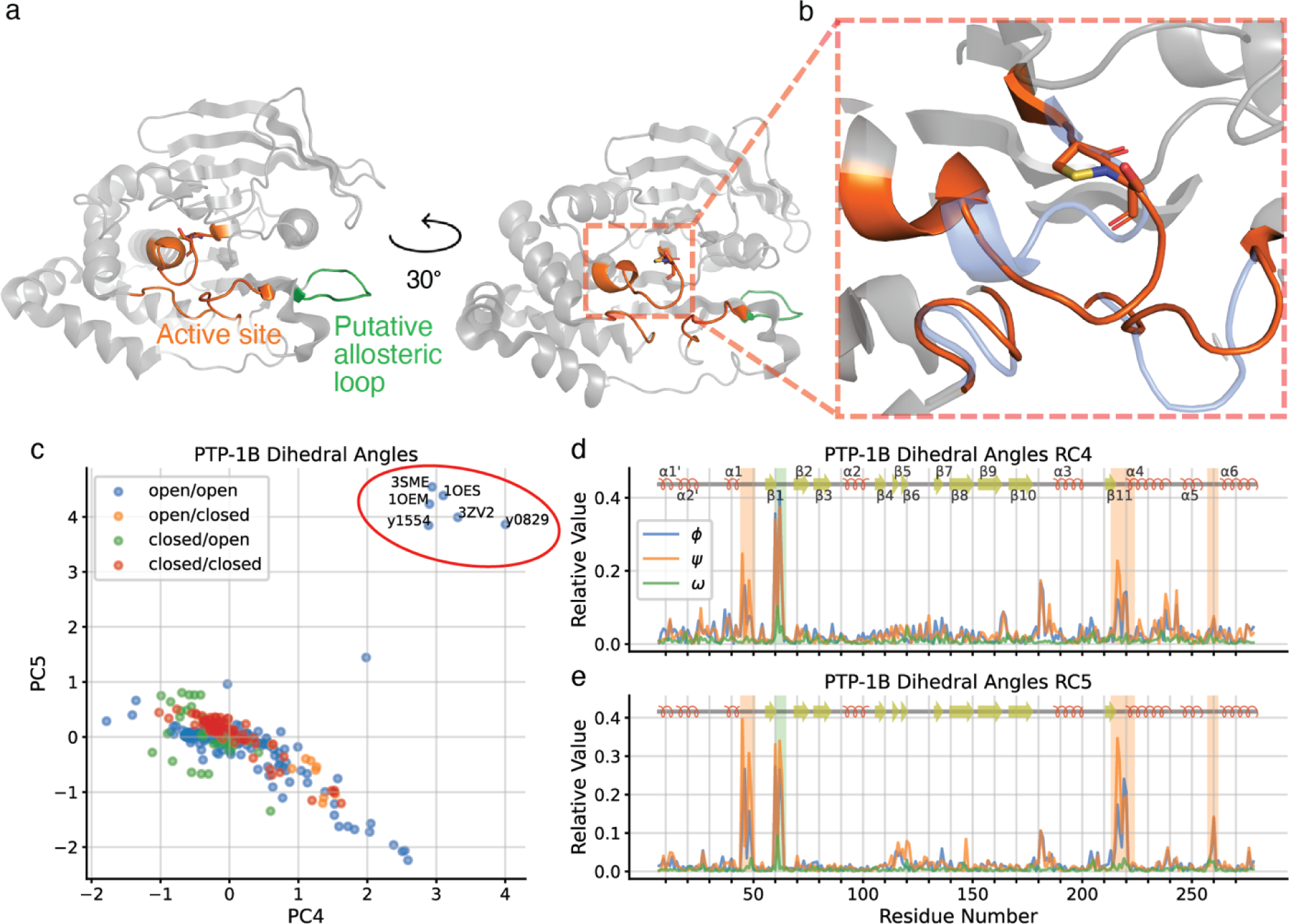
PTP-1B conformational change due to oxidation states of Cys215. (a) Cartoon representations of oxidized PTP-1B conformation (1OES), highlighting active site loops (orange) and putative allosteric loop (green). (b) Cartoon representation of the oxidized PTP-1B active site conformation (1OES; orange), with sulphenyl-amide ring shown in sticks, and the reduced PTP-1B active site conformation (1SUG; blue) for comparison. (c) PTP-1B conformational landscape by dihedral angles by PC4 and PC5; structures showing oxidized PTP-1B conformation as in panels (a) and (b) are circled in red. (d) Residue contributions to PC4, with active site loops in orange box and putative allosteric loop (residues 59-66) in green box (coloring matches panel (a)). (e) Residue contributions to PC5, with loop coloring as in panel (d).

In the analysis of these oxidized structures, we further noticed a strong signal from a region of PTP-1B distant from the active site and distinct from the L16 loop (green shaded box in Figure 3d,e). This spike in signal corresponds to a short loop including residues 59-66. Intriguingly, this loop is near Ser50 and contains Tyr66, two known phosphorylation sites of PTP-1B^69,70^. Furthermore, a computational analysis of PTP-1B by CryptoSite^71^ indicated that this loop is directly adjacent to a cryptic binding site capable of accommodating a small molecule. These observations point to a potential regulatory role of this loop in PTP-1B and perhaps a more direct role in the regulation of oxidized PTP-1B. Speculatively, recent work has shown that the E3-ligase Cullin1 is known to interact with oxidized (sulfonated Cys215) PTP-1B, but the mechanism of this molecular recognition is unclear. The putative coupling suggested by our analysis implies that oxidation of Cys215 triggers concerted motions in this loop, which may allow for recognition and ubiquitination by Cullin1.

### Drug fragment screen structures recapitulate the PTP-1B conformational landscape

Could structures from only the PTP-1B crystallographic drug fragment screen^36^ suffice to infer the same conformational landscape as the complete PTP-1B dataset or the (non-screen) PTP-1B structures deposited in the PDB (“PDB-only”)? To address this question, we again used the dihedral angle representation to map the conformational landscape of PTP-1B based solely on either the fragment screen or the PDB structures (Fig. 4). We first quantified the similarity of the fragment screen-only dataset and the PDB-only dataset using matching and coverage scores^72,73^. The matching score reports on how similar the datasets are by RMSD (root-mean-square deviation) and the score ranges from 0 (each structure has an identical match in the other dataset) to infinity. The coverage score reports on the relative diversity between the datasets and ranges between 0 and 1. Because these scores compare individual structures between datasets, comparing either the fragment screen-only or the PDB-only datasets to the complete dataset would yield perfect scores (matching score of 0 and coverage score of 1) because they contain the same structures, so we compared the fragment screen-only dataset and PDB-only dataset. We calculated the matching score to be 0.493 Å and the coverage score to be 0.963 with an RMSD similarity cutoff of 1.0 Å, which indicated that the fragment screen-only dataset resembles the PDB-only structures both in terms of containing similar (“matching”) structures and in the overall coverage of the conformational landscape.

**Figure 4:**
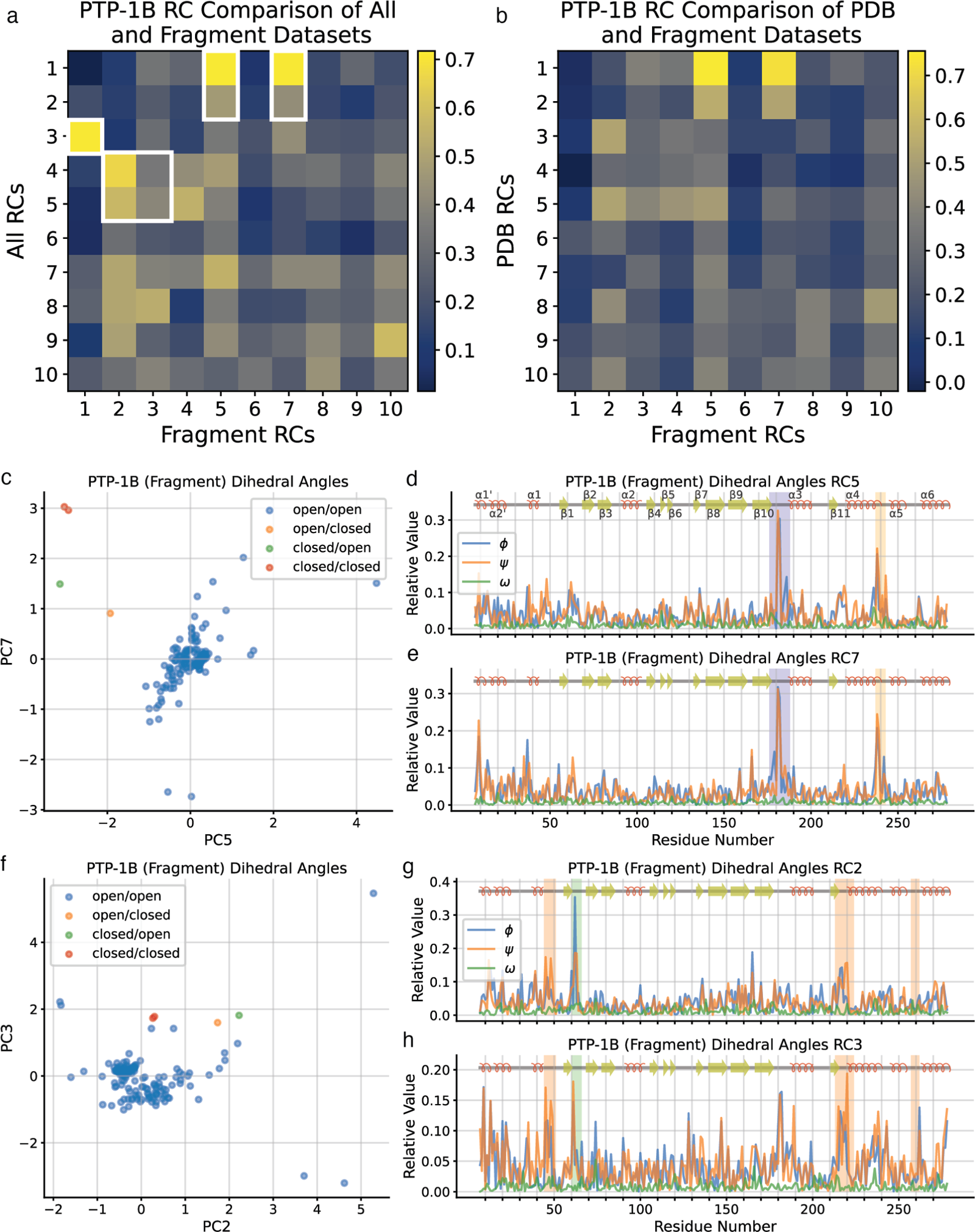
COLAV analysis of the PTP-1B crystallographic drug fragment screen recapitulates key aspects of the conformational landscape. (a, b) Correlation coefficient matrix comparing residue contributions (RCs) of (a) the complete PTP-1B dataset to those of the fragment screen-only PTP-1B dataset, and (b) the PDB-only PTP-1B dataset to those of the fragment screen-only PTP-1B dataset. Correlations discussed in the text are highlighted using white edges. (c) Fragment screen PTP-1B conformational landscape by dihedral angles, emphasizing similarities of PC5 and PC7 with PC1 and PC2 of the complete PTP-1B conformational landscape. (d) Residue contributions to PC5, with WPD loop in purple box and L16 loop in yellow box. (e) Residue contributions to PC7, with coloring as in panel (d). (f) Fragment screen PTP-1B conformational landscape by dihedral angles, emphasizing similarities of PC2 and PC4 with PC4 and PC5 of the complete PTP-1B conformational landscape. (g, h) Residue contributions to (g) PC2 and (h) PC4, with active site loops in orange box and putative allosteric loop in dark blue box.

To determine the relationship between the inferred conformational landscapes more carefully, we compared RCs for PCs from each dataset by calculating correlation coefficients. We found that most key RCs from the complete dataset were also clearly identifiable from the fragment screen-only dataset (Fig. 4a). We found similar results when we compared RCs of the fragment screen-only dataset and the PDB-only dataset (Fig. 4b). This mapping suggests similar structural interpretations for the complete, fragment screen-only, and PDB-only datasets. Indeed, the fifth and seventh fragment screen RCs resemble the first and second RCs of the complete dataset, again indicating a coupling between the WPD loop and L16 loop (Fig. 4c-e), albeit with different proportions of the major states. We note that since refinement of partial-occupancy states, typical for drug fragment screens, tends to be biased towards the unbound state, closed-loop conformations are likely underreported. Effects of catalytic cysteine oxidation were more prominent in the drug fragment screen than in the whole dataset, as observed by Keedy *et al*.^36^, such that the second and third fragment screen RCs correlated well with the fourth and fifth RCs of the complete dataset. As discussed above, the fourth and fifth RCs of the complete dataset report on active site deformation due to Cys-215 oxidation (Fig. 4f-h). We note that the first PC of the fragment-only dataset partially reports on a coupling between the L16 loop and the K loop, another active-site loop, that receives little weight in the PDB-only dataset (Fig. S5b). These comparisons show that the PTP-1B fragment screen conformational landscape matches that of the complete PTP-1B dataset, albeit with a different order of the PCs. This reordering reflects the relative prevalence of the different conformations in the fragment screen dataset.

### Continuous motions in the SARS-CoV-2 linker may be coupled to distant surface sites

We next applied the representations implemented in COLAV and PCA to the SARS-CoV-2 main/3CL protease (MPro). MPro is a component of a polyprotein translated from the positive-sense SARS-CoV-2 RNA genome. Through its protease activity, MPro cleaves itself and other functional proteins from this polyprotein, making MPro essential for viral replication^74^.

Consequently, MPro is a validated drug target for coronavirus disease caused by SARS-CoV-2 infection (COVID-19). The protein consists of three subdomains: domains I and II form a β-barrel catalytic core, and domain III forms an α-helical bundle unit that facilitates MPro obligate homodimerization (Fig. 5a,b)^75,76^. MPro is the subject of an intense research effort, with several crystallographic drug fragment screens and many other structural studies capturing the homodimer bound to a variety of ligands^37–41^. We analyzed 1,830 structures from these fragment screens and 1,015 other structures deposited in the PDB to determine the MPro conformational landscape by PCA.

**Figure 5:**
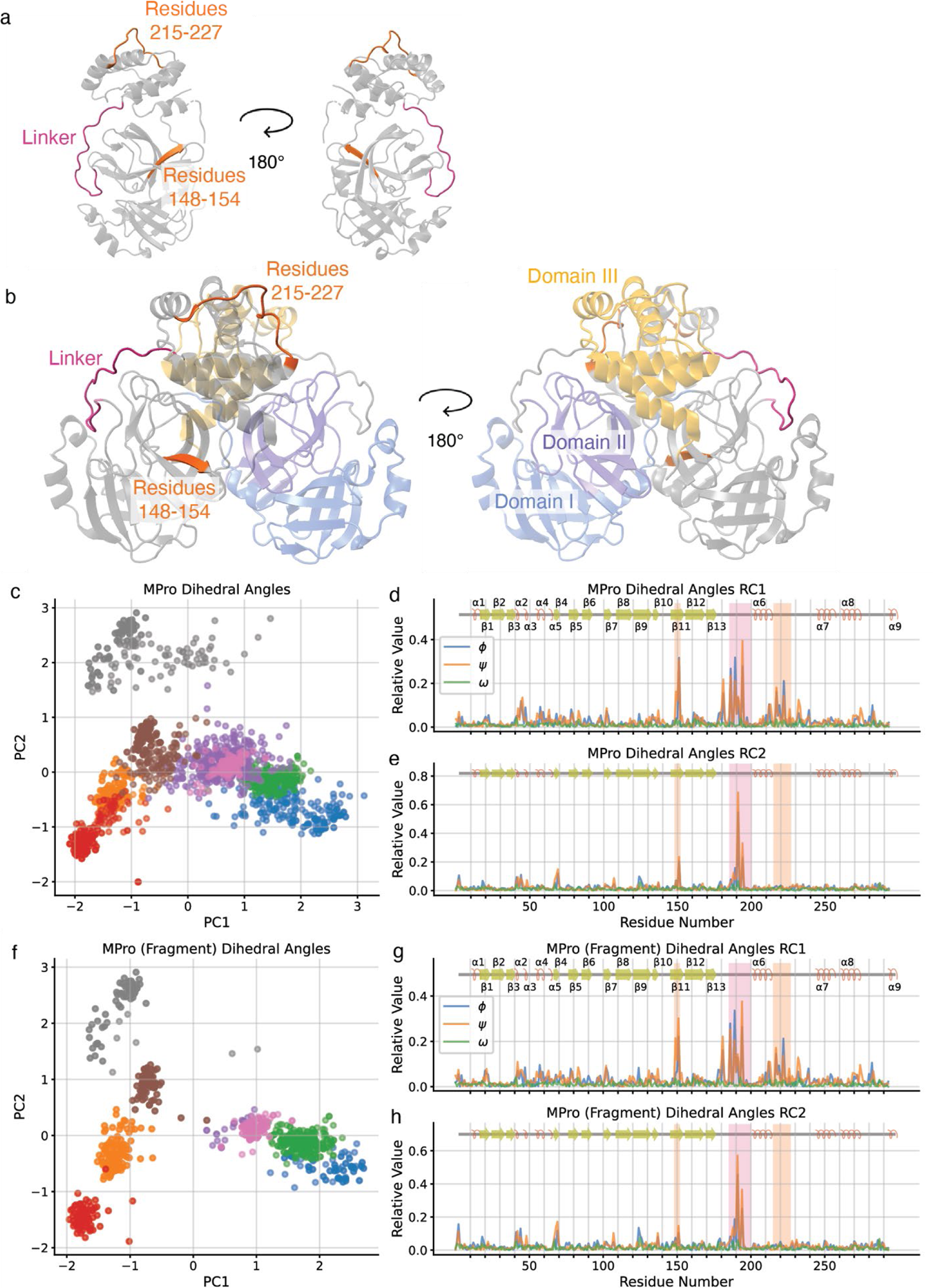
The MPro complete dataset and fragment-screen-only dataset generate similar conformational landscapes. (a) Cartoon representations of single MPro protomer (7AR5), highlighting linker (residues 185-200) in magenta and putative allosteric regions (residues 148-152 and 215-227) in orange. (b) Cartoon representations of MPro homodimer (7AR5), highlighting subdomain I in blue, subdomain II in purple, and subdomain III in yellow on protomer 1 and linker and putative allosteric regions colored as in (a). (c) MPro conformational landscape by dihedral angles. (d) Residue contributions to PC1, with linker in magenta box and putative allosteric loops in orange box (coloring matches panels (a) and (b)). (e) Residue contributions to PC2, with loop coloring as in panel (d). (f) Fragment screen MPro conformational landscape by dihedral angles. (g, h) Residue contributions to fragment screen (g) PC1 and (h) PC2, with loop coloring as in panel (d).

In contrast to PTP-1B, the MPro conformational landscape is dominated by a continuous band of structures along PC1 rather than by distinct clusters (Fig. 5c); along PC2, there is a distinct cluster of structures. We cautiously interpreted this to mean that the most common motions in MPro are continuous in the protein: the most flexible regions of the protein do not take on distinct, individual states. However, structures that are related in our conformational landscape, a reduced-dimensional space, may be more dissimilar in the higher-dimensional space considering all dihedral angles. To test this interpretation, we determined similar groups of MPro structures using the *k*-means algorithm (*k* = 8) for the full high-dimensional dihedral angle representation of each structure, yielding groups that are similar in the high-dimensional space. This proximity is well preserved in the low-dimensional space of the first two principal components (Fig. 5c). As for PTP-1B, PCA determined similar results for the three structural representations according to the *k*-means groups (Fig. S6; coloring of the structures matches between panels; t-SNE and UMAP analysis in Figure S6). From these analyses, we concluded that the dominant concerted motion in MPro is a gradual deformation.

To further investigate the motions of MPro and its correlated regions, we examined the residue contributions, again focusing on the dihedral angle representation. We interpreted the RCs corresponding to PC1 and PC2, respectively accounting for 14.4% and 7.0% of the total variance, as indicative of motion in the linker between domains II and III (Fig. 5d,e). Molecular dynamics simulations and ensemble refinement of MPro structures have shown that this region of the protein is flexible^42,77^. In addition, the motion corresponding to the first PC indicates that this linker is correlated with residues 148-154 and residues 215-227 (Fig. 5d,e). These regions are located approximately 20 Å and 30 Å away from the linker, respectively, in both a single protomer and the homodimer (Fig. 5a,b), indicating an allosteric coupling between these regions. Because the linker abuts the MPro active site, these regions may be suitable targets for drug design.

Next, we asked again whether the drug fragment screen recapitulates the conformational landscape inferred from either the complete MPro dataset or the non-fragment screen (“PDB-only”) dataset, as we did for PTP-1B above. We similarly find that the fragment screen-only dataset is nearly as conformationally diverse as the PDB-only dataset, with a coverage score of 0.925 using a RMSD threshold of 1.0 Å; a matching score of 0.466 Å shows that the structures of the fragment screen-only dataset closely match those of the PDB-only dataset. Likewise, we similarly find that the residue contributions to the different PCs have close matches between the fragment screen-only dataset and the whole dataset or the PDB-only dataset (Fig. S7). Therefore, as in PTP-1B, COLAV analysis of the MPro crystallographic drug fragment screen mapped the MPro conformational landscape efficiently and thoroughly.

We have found that conformational landscapes inferred from drug fragment screens alone recapitulate the main features of the conformational landscapes that can be inferred from larger ensembles of structures present in the PDB, often including deliberately designed mutants or targeted ligands. The stronger correspondence found for MPro (Figure 5) than for PTP-1B (compare Figure 4 to Figures 1-3) suggests that the sheer number of fragment-bound structures is an important parameter. To test this idea, we generated random samples from the MPro drug fragment screening dataset without replacement. We then compared the inferred conformational landscapes (based on dihedral angles) to the complete dataset by calculating correlation coefficients between RCs (Fig. S7). Compared to the complete dataset, we found that a reduced dataset of 135 structures was sufficient to broadly capture the top 5 RCs of the complete dataset (Fig. S7e). Most of the top 10 RCs were strongly recapitulated in the reduced datasets of 270 and 540 structures (Fig. S7c, d), matching the visual appraisal that the inferred conformational landscape looks like that of the complete dataset.

### Ordering protein structures by PC score exposes potential transition pathways

The PCA results showed several apparent conformational transitions in both PTP-1B and MPro. To examine these transitions more closely, we used the PC scores to order the structures of either PTP-1B or MPro for both PC1 and PC2 using the dihedral angle representation (Fig. 6). Doing so with PC1 for PTP-1B showed a distinct transition of the WPD loop between the open and closed state (Fig. 6a), while the same for PC2 described the transition of the L16 loop from a closed to open state (Fig. 6b). For MPro, the transitions between most conformations for PC1 and PC2 are more subtle (Fig. 6c, d), except for a distinct transition between MPro conformations in the linker along PC2 (Fig. 6d). Ordering structures by PC scores is especially informative when analyzing structures from crystallographic drug fragment screens, as conformations can be paired with the fragment ligands that stabilize them. Those fragment ligands that stabilize particular conformations of the target protein are then readily identifiable as the basis for targeted rational drug design.

**Figure 6:**
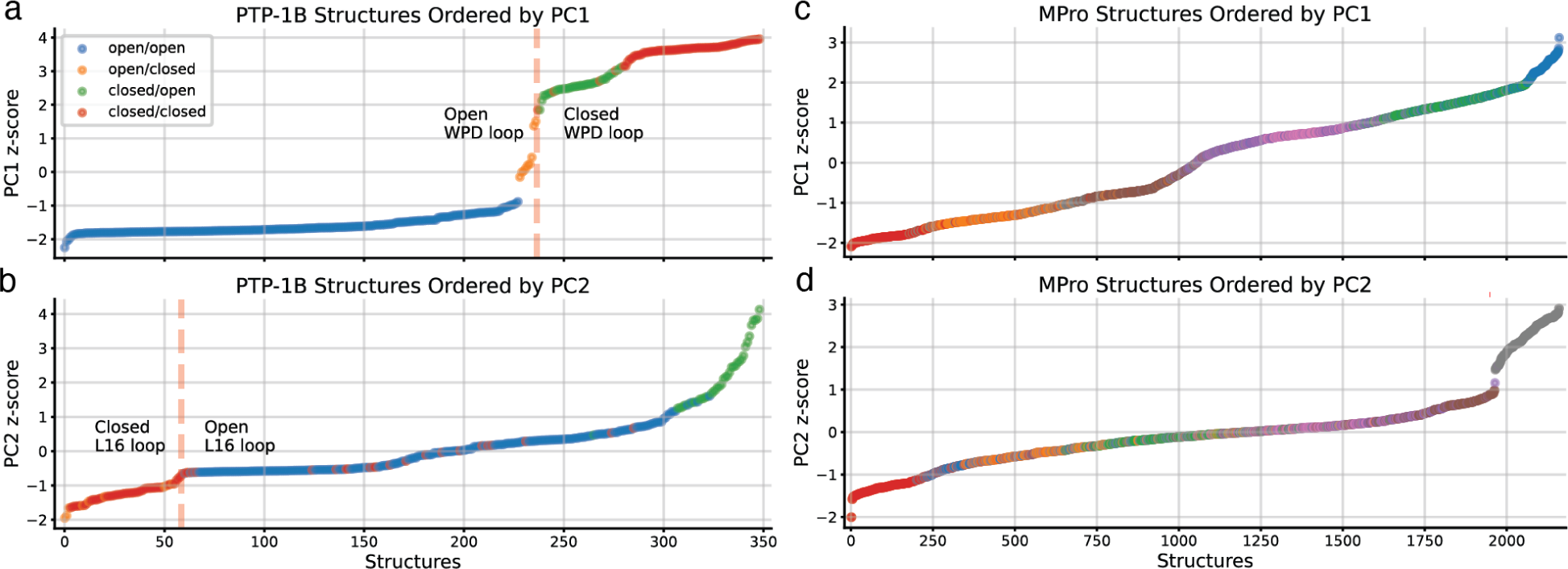
Ordering structures of PTP-1B and MPro by PC scores marks distinct conformational transitions. (a) PTP-1B structures ordered by dihedral angle score along PC1, with the transition from open WPD loop to closed WPD loop highlighted. (b) PTP-1B structures ordered by dihedral angle score along PC2, with the transition from closed L16 loop to open L16 loop highlighted. (c) MPro structures ordered by dihedral angle score along PC1. (d) MPro structures ordered by dihedral angle score along PC2. Coloring of datasets for both proteins matches preceding figures.

## Conclusions

Crystallographic drug fragment screens provide rich data, not only concerning the binding sites of fragments on drug targets but also on how protein conformations change in response to such binding. In this respect, drug fragment screens approximate an ideal experiment in which the structure of a protein is determined in the presence of each of many random perturbations. We introduced an open-source software package, COLAV, to facilitate inference of empirical conformational landscapes from such drug fragment screening data using three different representations of conformational change. We find that the results are insensitive to the choice of representation and largely robust under the choice of method for dimension reduction, indicating that the discovered conformational clustering is intrinsic to the conformational ensembles studied. Moreover, we found that the conformational landscapes determined this way resemble those inferred from the larger universe of previously determined structures and that the correspondence improves with the number of fragment-bound structures. Altogether, these findings lay the foundation for the systematic use of crystallographic drug fragment screens to map the accessible states of proteins of interest and a roadmap for steering proteins toward desirable conformations. The tools introduced in COLAV are general and may perform equally well for other protein structural ensembles, as the highly constrained nature of protein dynamics will leave its fingerprints on any such dataset.

## Author Contributions

AAS, MAK, and DRH conceived the approach. AAS developed the code and performed the analysis with feedback from MAK and DRH. All authors contributed to the manuscript.

## Acknowledgement

We thank Dr. Daniel Keedy and Dr. Helen Ginn, and members of the Hekstra lab for fruitful discussions. We thank Dennis Brookner for assistance in making COLAV available as a package on https://github.com/Hekstra-Lab/colav and PyPI.

## Data and Code Availability

All data and code used in this study to generate the figures can be found at https://github.com/Hekstra-Lab/colav. Figures were prepared using PyMOL v2.5.4, available from Schrödinger, LLC.

## Supplementary Figures and Table

**Figure S1:**
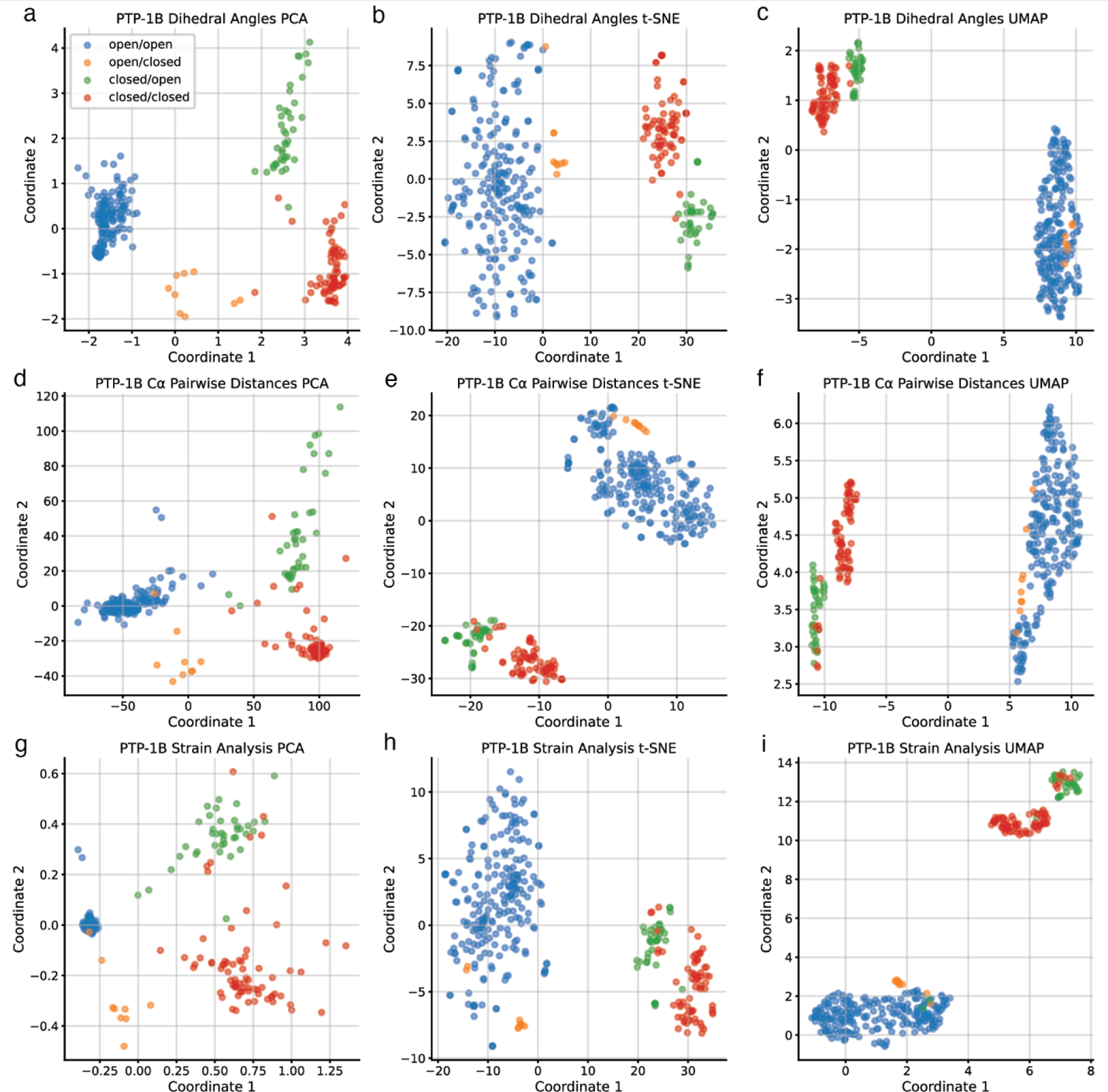
Comparison of PCA, t-SNE, and UMAP applied to all three structural representations of PTP-1B colored by conformation. (a-c) PTP-1B conformational landscape based on dihedral angles with dimensionality reduction by (a) PCA, (b) t-SNE, and (c) UMAP. (d-f) PTP-1B conformational landscape based on Cα pairwise distances, analyzed using (d) PCA, (e) t-SNE, and (f) UMAP. (g-j) PTP-1B conformational landscape based on strain analysis and (g) PCA, (h) t-SNE, and (i) UMAP. Coloring of structures is consistent among all panels.

**Figure S2:**
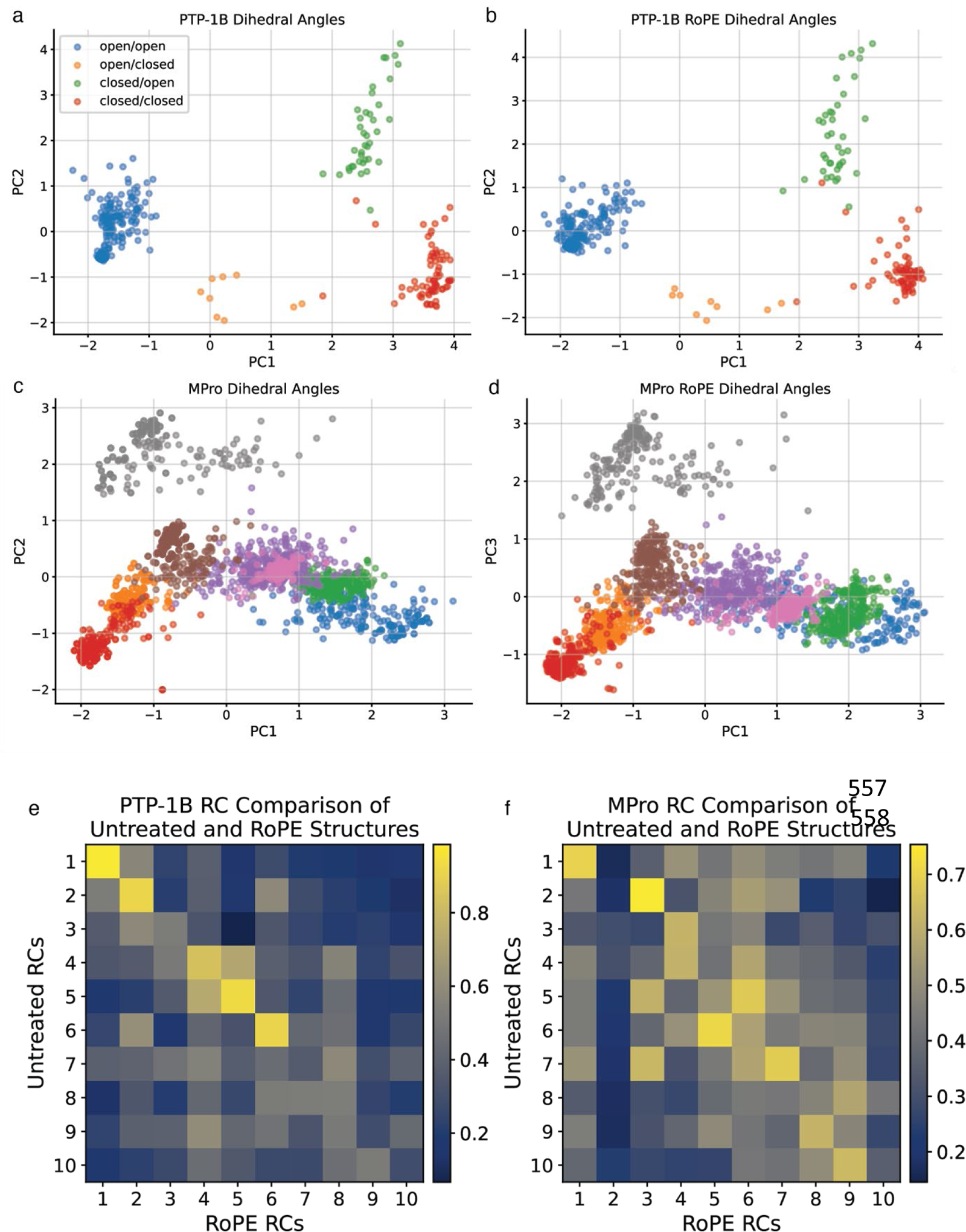
Dihedral angles with and without idealization by RoPE reveal similar conformational landscapes. (a) PTP-1B conformational landscape by dihedral angles calculated by COLAV. (b) PTP-1B conformational landscape by dihedral angles idealized by RoPE. (c) MPro conformational landscape by dihedral angles calculated by COLAV. (d) MPro conformational landscape by dihedral angles calculated by RoPE. (e) Correlation coefficient matrix comparing PTP-1B RCs of the untreated (COLAV) dihedral angles and RoPE dihedral angles. (f) Correlation coefficient matrix comparing MPro RCs of the untreated (COLAV) dihedral angles and RoPE dihedral angles.

**Figure S3:**
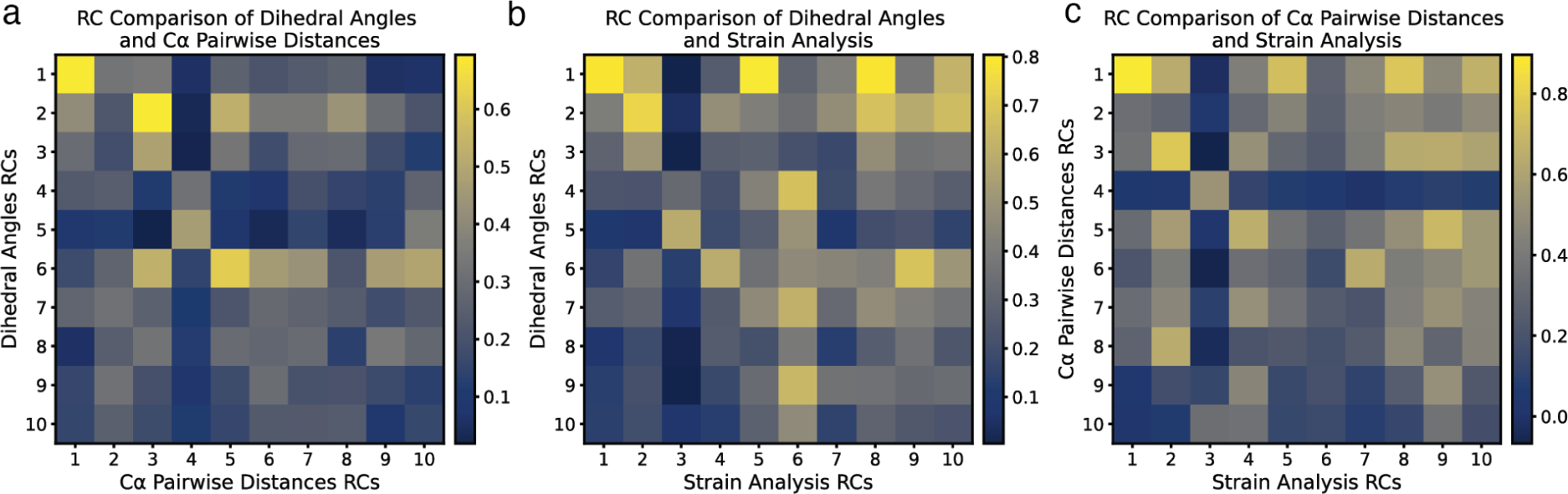
Comparison of residue contributions for structural representations of PTP-1B. Correlation coefficients comparing PTP-1B residue contributions (RCs) for (a) dihedral angles and Cα pairwise distances, (b) dihedral angles and strain, and (c) Cα pairwise distances and strain.

**Figure S4:**
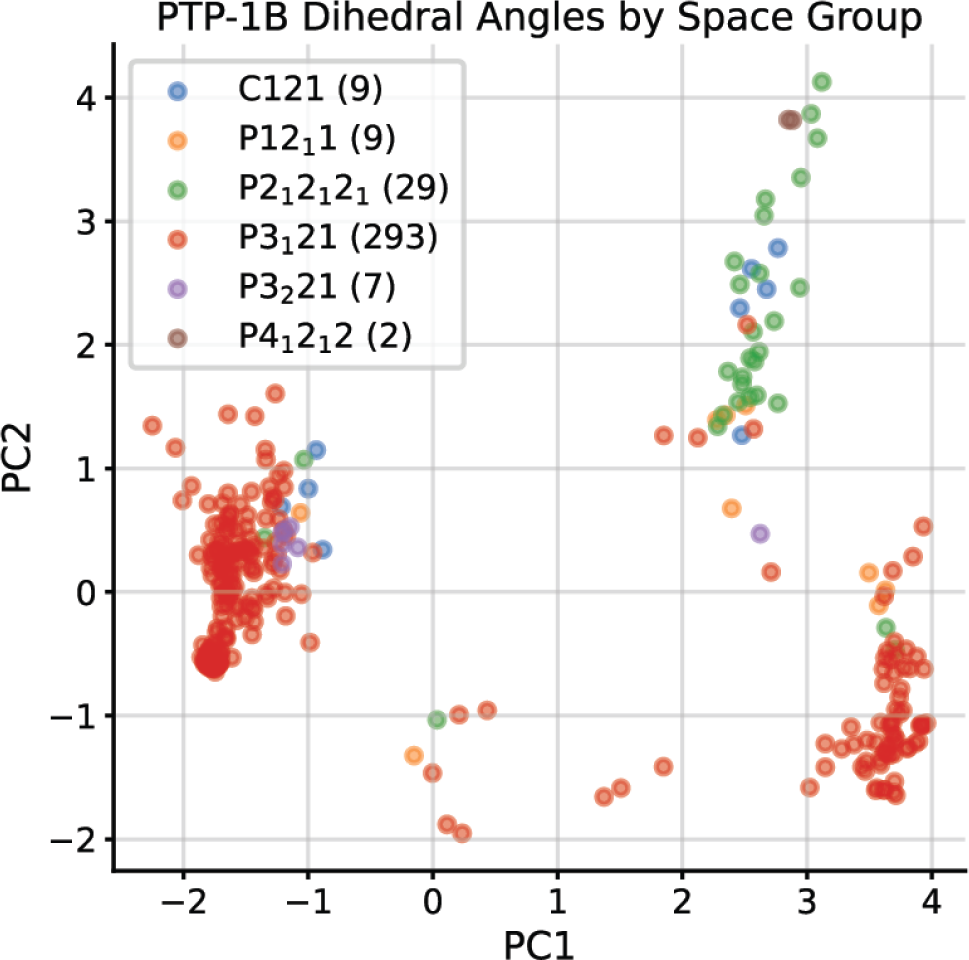
PTP-1B conformations are found across space groups. Distribution of structures of PTP-1B after PCA of their dihedral angles. Structures are colored by the space group of their crystal forms.

**Figure S5:**
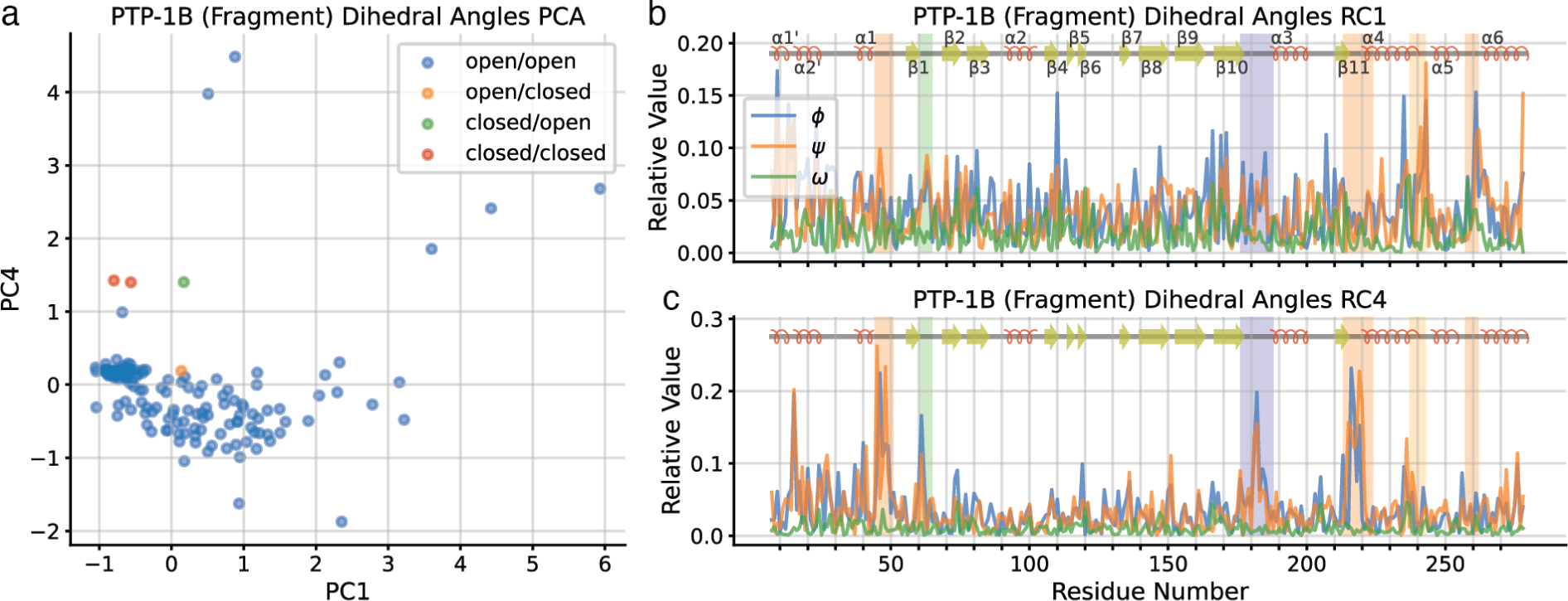
Additional dimensions of the PTP-1B conformational landscape inferred from crystallographic drug fragment screen structures. (a) Fragment screen PTP-1B conformational landscape by dihedral angles, using PC1 and PC4. (b) Residue contributions to PC1, with active site loops in orange box, putative allosteric loop in dark blue box, WPD loop in purple box, and L16 loop in yellow box. (c) Residue contributions to PC4, with coloring as in panel (b).

**Figure S6:**
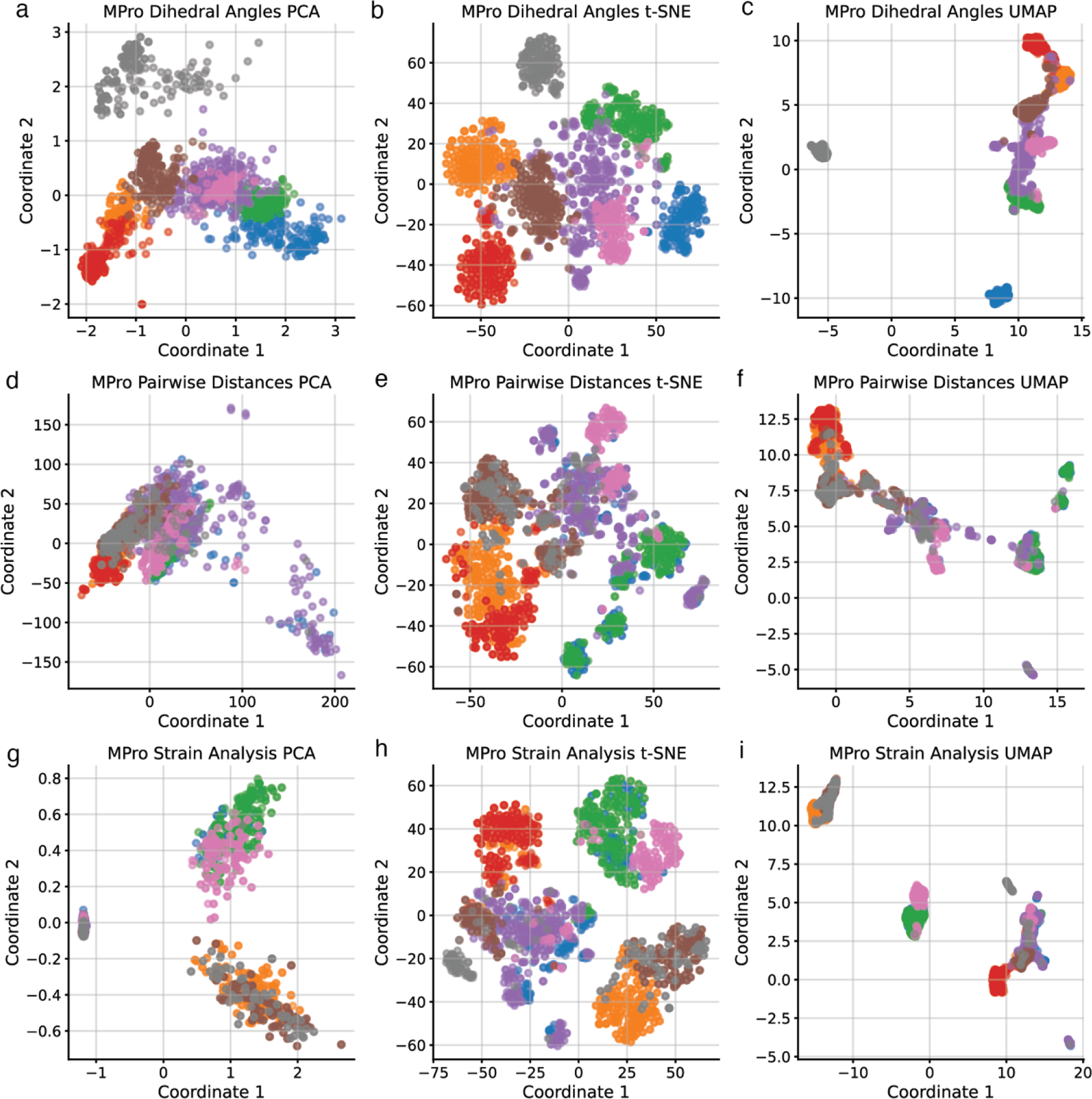
Comparison of PCA, t-SNE, and UMAP applied to all three structural representations of MPro. (a-c) MPro conformational landscape based on dihedral angles with dimensionality reduction by (a) PCA, (b) t-SNE, and (c) UMAP. (d-f) MPro conformational landscape based on Cα pairwise distances, analyzed using (d) PCA, (e) t-SNE, and (f) UMAP. (g-j) MPro conformational landscape based on strain analysis and (g) PCA, (h) t-SNE, and (i) UMAP. Coloring of structures is consistent among all panels.

**Figure S7:**
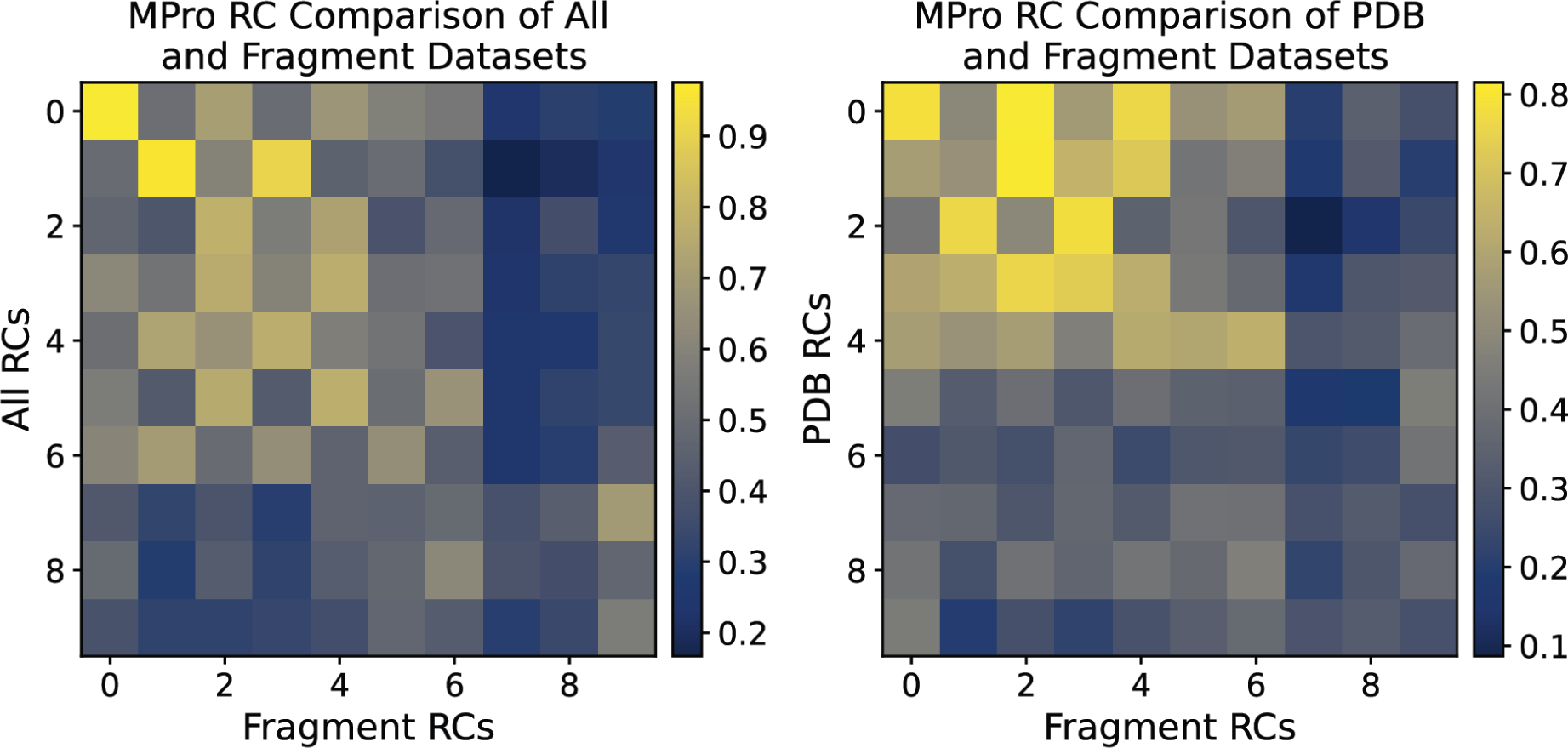
Comparison of dihedral angles residue contributions for MPro datasets. (a) Correlation coefficient matrix comparing RCs of the complete MPro dataset to those of the fragment screen-only MPro dataset. (b) Correlation coefficient matrix comparing RCs of the PDB-only MPro dataset to those of the fragment screen-only MPro dataset.

**Figure S8:**
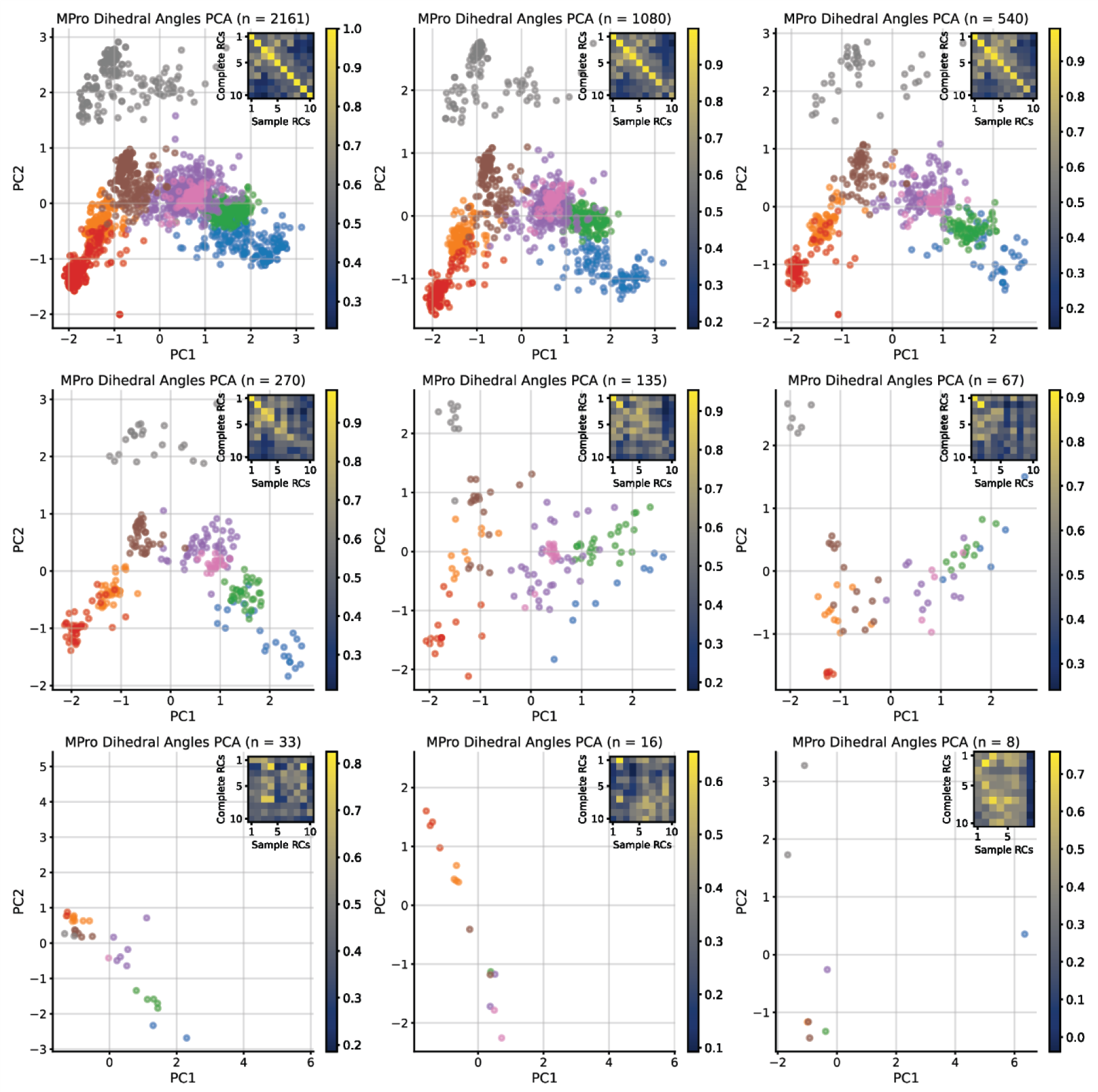
Effect of dataset size on the quality of inferred MPro conformational landscapes. MPro conformational landscape by dihedral angles were determined for the complete dataset and subsampled datasets of N = 2161 (a), 1080 (b), 540 (c), 270 (d), 135 (e), 67 (f), 33 (g), 16 (h) or 8 (i) structures. In each panel we include an inset of the correlation of the residue contributions inferred for the complete dataset and the sampled dataset.

**Table S1:**
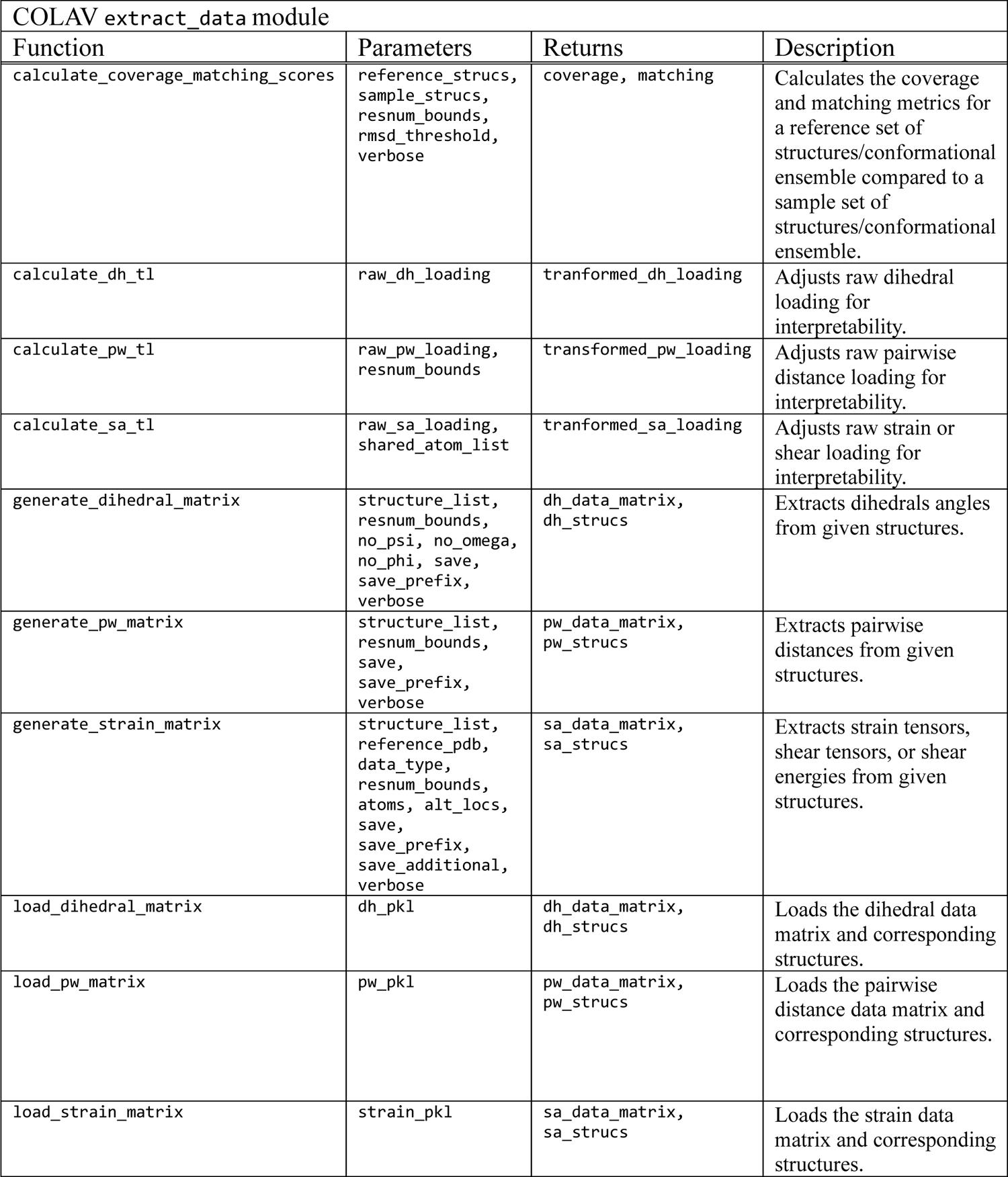
User-accessible COLAV functions for analyzing structural data. For a more complete description of the COLAV software package and its functionality, visit https://github.com/Hekstra-Lab/colav. Note that “transformed loadings” are referred to in the text as “residue contributions”.

